# Motor Hotspot Localization Based on Electroencephalography Using Convolutional Neural Network in Patients with Stroke

**DOI:** 10.1101/2024.03.06.583618

**Authors:** Ga-Young Choi, Jeong-Kweon Seo, Kyoung Tae Kim, Won Kee Chang, Nam-Jong Paik, Won-Seok Kim, Han-Jeong Hwang

**Affiliations:** Departmet of Bioengineering, McGill University, Montreal, Quebec, Canada; Departmet of Electronics and Information Engineering, Korea University, Sejong 30019, Republic of Korea; Institute of Data Science, Korea University, Seoul, South Korea; Department of Rehabilitation Medicine, Keimyung University School of Medicine, Keimyung University Dongsan Hospital, Daegu, Republic of Korea; Department of Rehabilitation Medicine, Seoul National University College of Medicine, Seoul National University Bundang Hospital, Seongnam-si 13620, Republic of Korea; Interdisciplinary Graduate Program for Artificial Intelligence Smart Convergence Technology, Korea University, Sejong 30019, South Korea

**Keywords:** Deep learning, Motor hotspot, Electroencephalography, Neurorehabilitation

## Abstract

**Background:** Although transcranial magnetic stimulation (TMS) is the optimal tool for identifying individual motor hotspots for transcranial electrical stimulation (tES), it requires a cumbersome procedure in which patients must visit the hospital each time and rely on expert judgment to determine the motor hotspot. Therefore, in previous study, we proposed electroencephalography (EEG)-based machine learning approach to automatically identify individual motor hotspots. In this study, we proposed an advanced EEG-based motor hotspot identification algorithm using a deep learning model and assessed its clinical feasibility and benefits by applying it to stroke patient EEGs.

**Methods:** EEG data were measured from thirty subjects as they performed a simple hand movement task. We utilized the five types of input data depending on the processing levels to assess the signal processing capability of our proposed deep learning model. The motor hotspot locations were estimated using a two-dimensional convolutional neural network (CNN) model. The error distance between the 3D coordinate information of the individual motor hotspots identified by the TMS (ground truth) and EEGs was calculated using the Euclidean distance. Additionally, we confirmed the clinical benefits of our proposed deep-learning algorithm by applying the EEG of stroke patients.

**Results:** A mean error distance between the motor hotspot locations identified by TMS and our approach was 2.34 ± 0.19 mm when using raw data from only 9 channels around the motor area. When it was tested on stroke patients, the mean error distance was 1.77 ± 0.15 mm using only 5 channels around the motor area.

**Conclusion:** We have demonstrated that an EEG-based deep learning approach can effectively identify the individual motor hotspots. Moreover, we validated the clinical benefits of our algorithm by successfully implementing it in stroke patients. Our algorithm can be used as an alternative to TMS for identifying motor hotspots and maximizing rehabilitation effectiveness.

## Introduction

According to a World Health Organization (WHO) report, stroke-related mortality reached 6.6 million cases in 2019, ranking third as the cause of death worldwide [1]. Stroke primarily leads to motor impairments, which not only limit daily activities but also impose significant restrictions on social engagements. It has a significant impact on the quality of life for affected individuals, including both patients and caregivers, and can lead to chronic health problems. Therefore, it is necessary to thoroughly comprehend these issues and develop effective rehabilitation programs. One of the efficient neurorehabilitation methods is transcranial electrical stimulation (tES), which is a non-invasive brain stimulation technique to modulate neuronal excitability, including transcranial direct current stimulation (tDCS), and transcranial alternating current stimulation (tACS) [2–7]. Numerous studies have shown that tES stimulation is widely recognized as an effective treatment for improving motor function after a stroke [8], particularly in enhancing hand movement skills [9–13].

Standard tES typically comprises a single anode and cathode electrode. To enhance hand motor function in stroke patients, the anode electrode is positioned on the primary motor cortex (M1) in the contralateral hemisphere of the paralyzed hand. The cathode electrode is positioned either at the ipsilateral M1 site or at the supraorbital (SO) region of the same hand. For example, in order to enhance motor function in the right hand, the anode electrode is placed on the M1 site in the left hemisphere, while the cathode electrode is placed on either the M1 or SO region in the right hemisphere [14–17]. There are two approaches to identifying the precise brain region of hand movement within the M1 responsible. The first approach uses the international 10–20 system to estimate the location of the hand knob, which is the anatomical region that governs hand movements [15, 18–19]. The hand knobs are located in the central lobes of left and right hemispheres, referred to as C3 and C4 in the international 10–20 system, respectively. This approach is a quick and convenient method to identify brain regions responsible for rough hand movements. However, Kim et al. have shown that it is more effective in precisely stimulating a specific motor area, called a motor hotspot [14, 20–31]. The motor hotspot refers to the functional regions responsible for governing hand movements. The motor hotspot is typically located anteriorly and laterally in comparison to the hand knob, and its location can vary among individuals [28, 32]. To identify the motor hotspot, another stimulation device called transcranial magnetic stimulation (TMS) is required. An optimal target location for tES has been identified, where a maximum motor evoked potential (MEP) is generated in at least 50% of the TMS applications [33–36]. The TMS-based approach, commonly employed method for identifying individual motor hotspots, necessitates the involvement of experts and the utilization of specialized equipment, such as TMS and electromyography (EMG).

Therefore, in our previous study, we proposed a machine learning-based algorithm that used electroencephalography (EEG) to automatically identify individual motor hotspots, eliminating the need for TMS and EMG. By combining commercially available portable tES-EEG integrated devices (e.g., Starstim tES-EEG systems by Neuroelectrics, M×N-5, and M×N-9 HD-tES by Soterix Medical) with the developed algorithm, the accessibility of tES rehabilitation can be improved [37–38]. In other words, it has the potential to support home-based neurorehabilitation for patients with limited mobility by automatically identifying the individual motor hotspots based on EEGs. Consequently, the efficiency of motor rehabilitation can be improved by replacing expensive and bulky transcranial magnetic stimulation (TMS) equipment with less expensive electroencephalography (EEG) equipment. This not only simplifies the neurorehabilitation process but also provides economic benefits. However, the previous study has only validated the feasibility of the EEG-based motor hotspot identification approach on healthy individuals. Further research is required to verify the effectiveness of EEG patterns in stroke patients because the location of lesion and its severity can cause variations in the patterns [39]. Therefore, we propose a more robust and convenient algorithm for identifying EEG-based motor hotspots adapted to the individual characteristics of stroke patients. We also aimed to simplify the process of domain-knowledge-based feature engineering, which involves signal processing and feature extraction in machine learning, by utilizing a deep learning algorithm based on convolutional neural network (CNN). To do this, we initially assessed the identification performance of CNN model by testing its signal processing capability on EEG data from healthy subjects. We evaluated the model’s performance at various levels of handcrafted signal processing and then examined how performance changed by adjusting the number of channels and trials to assess the robustness of proposed model. Then, we ultimately validate its clinical benefits by applying EEG from stroke patients.

## Methods

### 1. Subjects

Thirty healthy subjects (10 females and 20 males; 25 ± 1.39 years; all right-handed) were recruited for this study. The participants had no prior psychiatric or neurological disorders that could affect the research. Before the experiment, participants were provided with information about the experimental procedure and were required to sign an informed consent form in order to participate in the study. Adequate reimbursement was provided for their participation in the experiment. The study protocol was approved by the Institutional Review Board (IRB) of Kumoh National Institute of Technology (No. 6250). The study was conducted in accordance with the Code of Ethics of the World Medical Association (Declaration of Helsinki).

### 2. Traditional Motor Hotspot Identification by TMS

Before conducting EEG measurements, we identified the motor hotspots in both hands for each subject by analyzing the TMS-induced MEPs of the first dorsal interosseous (FDI) muscle. We used Ag-AgCl disposable electrodes (actiChamp, Brain Products GmbH, Gilching, Germany) to measure the MEPs. We used single-pulse TMS on the contralateral motor cortex (REMED, Daejeon, Korea) to locate the motor hotspot. The motor hotspot was defined as the area where the maximum MEP of at least 50 μV was elicited in more than 5 out of 10 consecutive stimuli, using the individual’s minimum stimulation intensity [33–36]. To record the locations of motor hotspots, we used a digitizer (Polhemus Inc., Colchester, Vermont, USA) to mark the 3D coordinates based on the vertex (Cz) in the international 10–20 system. The coordinates provided served as the ground truth for comparing with the estimates obtained through our EEG-based motor hotspot identification approach.

### 3. EEG Recording

The sixty–three EEG electrodes were placed on the scalp according to the international 10– 20 system to measure EEG data. The ground and reference electrodes were attached to Fpz and FCz, respectively (Figure 1(a)). The EEG data were sampled at a rate of 1,000 Hz using a multi-channel EEG acquisition system (actiChamp, Brain Products GmbH, Gilching, Germany). The subjects performed a simple finger-tapping task to collect EEG data related to movement. They were instructed to press the spacebar with their index fingers whenever a red circle appeared in the center of a monitor (Figure 1(b)). Each trial consisted of a hand movement task followed by a relaxation period lasting between 3 and 7 s. It was repeated 30 times for each hand, with sufficient rest periods to prevent excessive fatigue whenever they wanted. EEG measurements were conducted independently using the index finger of the left and right hand, respectively. They were instructed to remain relaxed during the experiment and to avoid any unnecessary movements in order to prevent physiological artifacts. Prior to this experiment, the right hand was only used for two subjects because preliminary experiments were conducted on the first two subjects to verify the experimental paradigm. In addition, we excluded the EEG data of one subject for both hands and the data of the left hand for other three subjects due to high contamination of EEG data caused by physiological artifacts. Thus, 29 and 25 EEG datasets were used for the right and left hand, respectively, for data analysis.

**Figure 1.**
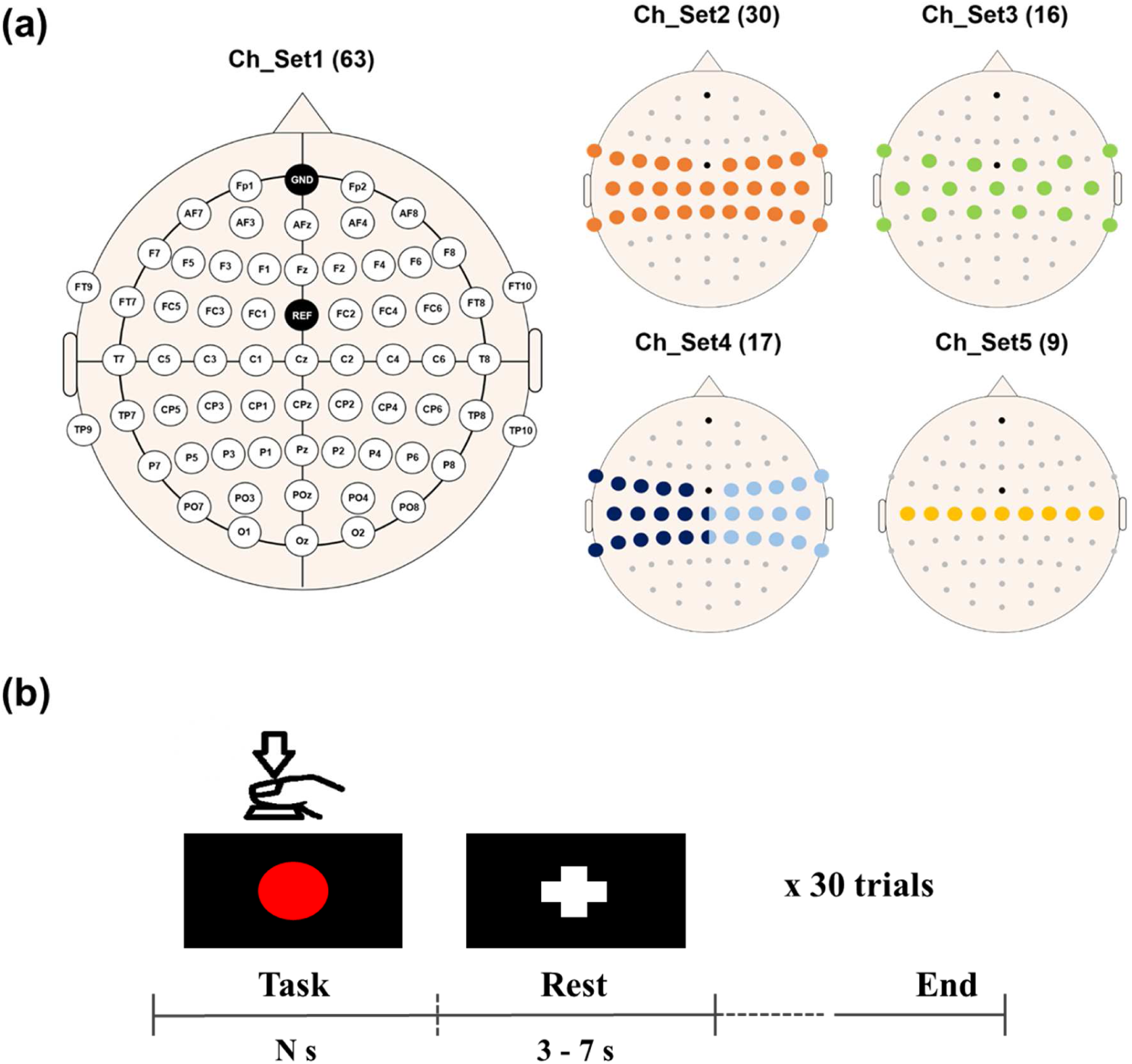
(a) Electrode attachment location used for EEG data in the healthy group. Five different channel sets are used for data analysis to determine the number of channels on the error distance in the motor hotspot location. Channels in different colors in Ch_Set4 represent those selected channels in the contralateral motor cortex for the data analysis of each hand. (b) Experimental paradigm. Each healthy subject presses a space bar whenever the red circle appears in the center of a monitor. The red circle remains until the subject presses the space bar. At the end of the task period, the fixation (‘+’) mark is displayed to indicate a rest period.

### 4. Data Analysis

#### 4.1 Pre-processing

We performed EEG data preprocessing using the EEGLAB toolbox based on MATLAB 2017b (MathWorks, Natick, MA, USA). The raw data were down-sampled to 200 Hz to simplify the calculation process. We sequentially applied common average reference (CAR) and band-pass filtering between 1 and 55 Hz using a zero-phase 3rd-order Butterworth filter. The filtered data underwent independent component analysis (ICA) to eliminate physiological artifacts. We excluded visually contaminated IC components. After preprocessing, we segmented the EEG data between -0.5 and 0.5 s based on the key press point for each trial. Power spectral densities (PSDs) of each EEG trial were estimated in the gamma frequency bands (30 – 50 Hz), which were identified as optimal features for identifying motor hotspots using EEG in our previous study [37]. The estimation was performed at a resolution of 1 Hz using the fast Fourier transform (FFT).

The extracted features, along with the 3D coordinates of motor hotspots identified by TMS-induced MEPs, were used as input for training the CNN model. To assess the signal processing capabilities of proposed CNN model, the input values were divided into five stages, using signal-processed input values at various levels. The first input, referred to as Input_1, consists of segmented time series EEG data ranging from -0.5 seconds to 0.5 seconds, based on the time point of key press. Input_2 consisted of time series EEG data that were re-referenced using CAR and segmented for the same period. Input_3 underwent sequential processing steps, including CAR, band-pass filtering from 1 to 55 Hz, and segmentation of the data within that period. Input_4 consists of time series EEG data that has undergone sequential processing steps, including CAR, band-pass filtering between 1 and 55 Hz, data segmentation, and ICA to remove physiological artifacts. Lastly, Input_5 consisted of PSD features that underwent sequential processing. This included CAR, band-pass filtering between 1 and 55 Hz, data segmentation, ICA, and FFT to extract the PSDs. The information is summarized in Table 1, and the format of the input data depends on the specific type of input. The model produced three-dimensional coordinates of the motor hotspot, estimated using EEG characteristics.

**Table 1.** Stages of input data for verifying the signal processing capability.

| Name | Contents | Format of input |
| --- | --- | --- |
| Input_1 | Raw | channel × time |
| Input_2 | Re-referencing (common average reference: CAR) | channel × time |
| Input_3 | CAR + band-pass filter (BPF) | channel × time |
| Input_4 | CAR + BPF + independent component analysis (ICA) | channel × time |
| Input_5 | CAR + BPF + ICA + fast Fourier transform | channel × PSDs |

#### 4.2 Two Dimensional-Convolutional Neural Network Model

Our proposed two-dimensional CNN (2D-CNN) model is implemented using the Keras library, an extension of TensorFlow. As depicted in Figure 2, the 2D-CNN model comprises 17 layers, which include 10 convolution layers, 4 pooling layers, and 3 fully connected layers [40–41]. In a convolutional layer, the kernel slides over the input to create the feature map. The pooling layer is used in layers 3, 6, 10, and 14 to decrease the dimension of the feature map. In this study, max-pooling was applied to the entire pooling layers, with the option of using two window strides in one step, while also incorporating zero-padding. The rectified linear unit (ReLU) was used as the activation function for all layers. This function extracts and transmits the final output signal to the next neuron. However, in the last fully connected layer, a linear activation function is used to regress the locations of the motor hotspots. The specifications of the proposed model for each layer, including kernel size, stride, etc., are summarized in Table 2. The proposed model was trained using the adaptive moment (ADAM) optimization algorithm to minimize the mean square error (MSE) loss function. The learning rate was set to 10^-3^ and the batch size for the training dataset was set to 10. For training the model, the number of iterations was set to 500 epochs, and early stopping based on validation loss was implemented to prevent overfitting. The patience parameter is set to allow training to be stopped after 20 additional verifications, based on the epoch point where the validation loss stops decreasing.

**Figure 2.**
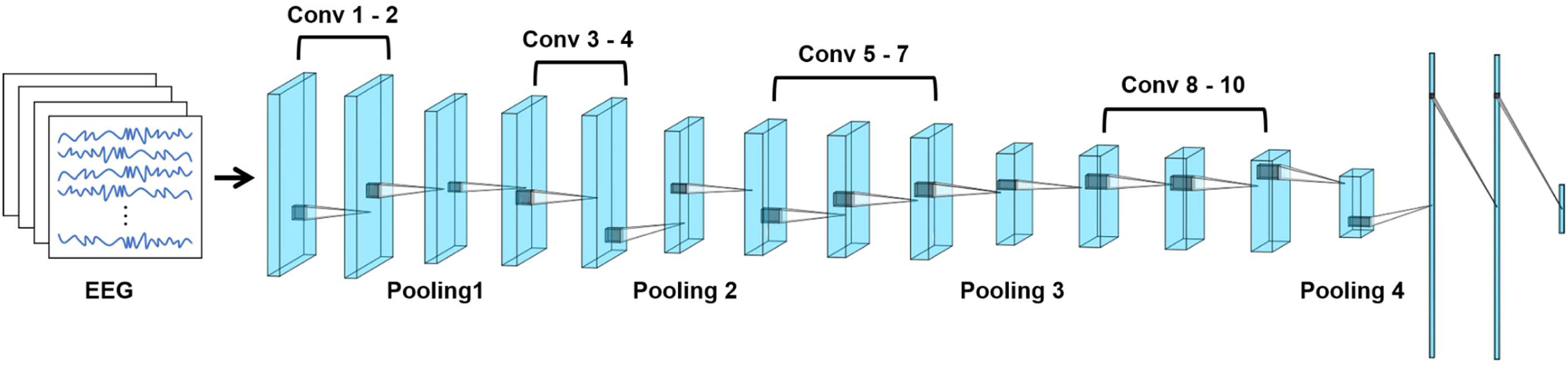
Schematic diagram of proposed two dimensional-convolutional neural network (2D-CNN) for identifying EEG-based motor hotspot location.

**Table 2.** Details of layer parameters of the proposed CNN model.

| Layer | Layer Name | Kernel | Number of Filters | Stride |
| --- | --- | --- | --- | --- |
| 0 | Input | - | - | - |
| 1 | Conv1 | 3 × 3 | 16 | 1 |
| 2 | Conv2 | 3 × 3 | 16 | 1 |
| 3 | Max-pooling1 | 2 × 2 | 16 | 2 |
| 4 | Conv3 | 3 × 3 | 32 | 1 |
| 5 | Conv4 | 3 × 3 | 32 | 1 |
| 6 | Max-pooling2 | 2 × 2 | 32 | 2 |
| 7 | Conv5 | 3 × 3 | 64 | 1 |
| 8 | Conv6 | 3 × 3 | 64 | 1 |
| 9 | Conv7 | 3 × 3 | 64 | 1 |
| 10 | Max-pooling3 | 2 × 2 | 64 | 2 |
| 11 | Conv8 | 3 × 3 | 128 | 1 |
| 12 | Conv9 | 3 × 3 | 128 | 1 |
| 13 | Conv10 | 3 × 3 | 128 | 1 |
| 14 | Max-pooling4 | 2 × 2 | 128 | 2 |
| 15 | Flatten | - | - | - |
| 16 | Dense1 | - | - | - |
| 17 | Dense2 | - | - | - |
| 18 | Dense3 | - | - | - |

The 2D-CNN model was trained and tested using trial-wise-based nested cross-validation to build a user-adapted model for understanding individual characteristics. Each subject’s data (30 trials) were independently used for model training (Figure 3) [37]. A 5×5-fold cross-validation was used in the outer loop of nested cross-validation. Each subject’s data were randomly divided into 80% for training and 20% for testing. In the inner loop, the 80% training dataset of outer loop was divided into a 4:1 ratio for training and validation data in order to determine the optimal hyperparameters. The prediction performance was evaluated using the test dataset from the outer loop. The evaluation process was repeated 25 times (5×5-fold) to ensure robustness and reliability. The performance of the final model was reported based on the averaged outputs from these iterations.

**Figure 3.**
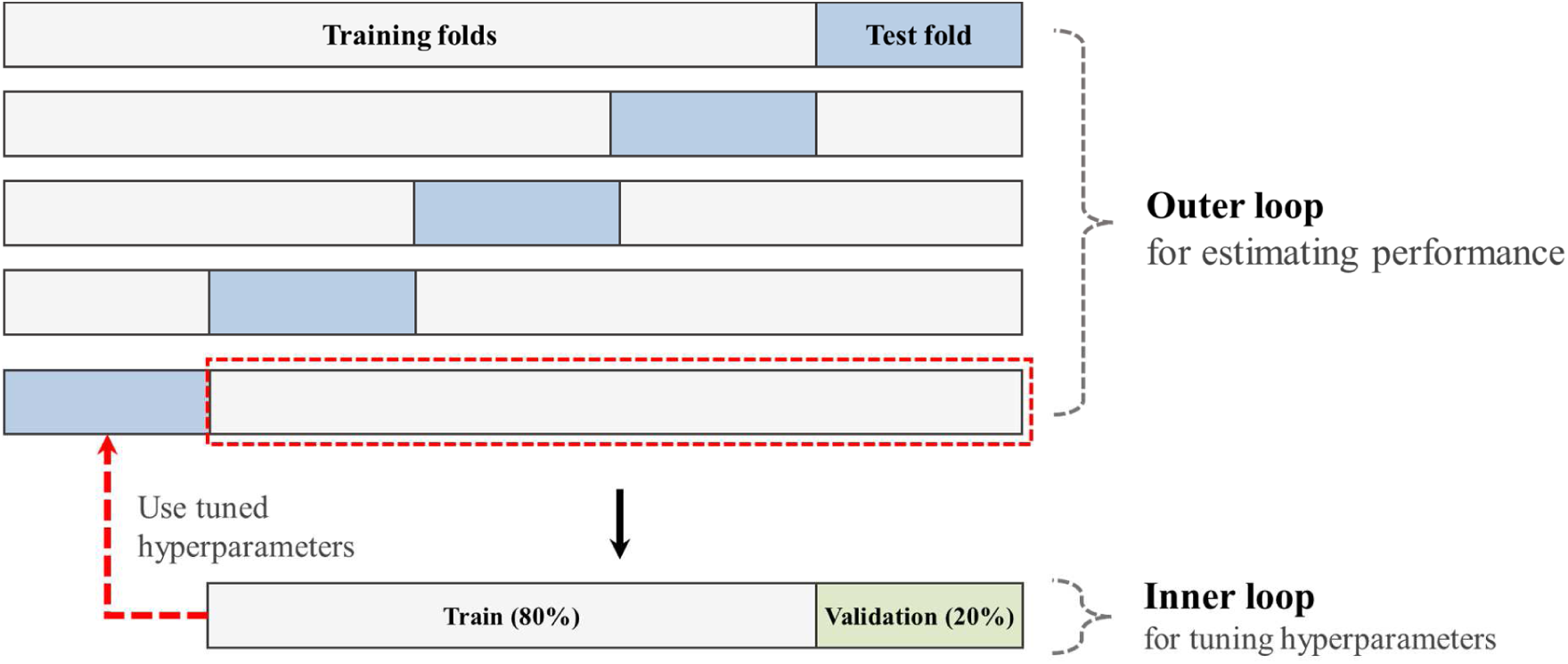
Scheme of the nested 5-fold cross-validation procedure.

### 5. Performance Verification

The error distance was quantitatively estimated by calculating the Euclidean distance between the 3D coordinates of the motor hotspot identified by TMS-induced MEP and the EEG-based deep learning approach. Additionally, the impact of the number of channels and trials on motor hotspot identification performance was investigated. This was done by gradually reducing the number of channels and trials, with a focus on the central area responsible for motor function (Figure 1(a)).

### 6. Verification of Clinical Benefits with Stroke

#### 6.1 Subjects

The stroke patients were enrolled in a department of rehabilitation in a tertiary hospital as subjects. Twenty–nine patients with acute stroke (5 females and 24 males; 63 ± 12 years; all right-handed except for one subject) were included in this study. We used the following criteria to enroll stroke patients: 1) patients aged 18 to 85 with impaired upper limb function; 2) confirmed ischemic or hemorrhagic stroke using a neuroimaging technique, such as CT or MRI; 3) patients with a Korean Mini-Mental State Exam (K-MMSE) score above 16 points; 4) patients capable of following instructions for clinical assessment and EEG study. Exclusion criteria included: (1) patients with traumatic brain injury or uncontrolled internal/surgical disease; (2) patients with other disorders, such as altered consciousness or psychiatric disorders; (3) pregnant patients; (4) patients with implanted pacemakers, cochlear implants, or a history of brain surgery. Participants were provided with information regarding the experimental procedure and signed informed consent forms to participate in the study. If the patient meets our criteria but is unable to provide consent due to a disability, the legal representative will sign on their behalf. The study protocol was approved by the Seoul National University Bundang Hospital IRB (registration No.: B-1912-580-005). The study was conducted in accordance with the Code of Ethics of the World Medical Association (Declaration of Helsinki).

#### 6.2 Experimental Protocol

The patients attached disposable Ag-AgCl electrodes (Synergy EMG/EP system; Oxford Instruments Medical Ltd., Surrey, UK) to the FDI muscle and applied a single TMS pulse (MagPro X100; MagVenture, Farum, Denmark) to establish the ground truth for each hand, following the procedure outlined in **’2. Traditional Motor Hotspot Identification by TMS’** prior to the experiment. If only unilateral MEP was identified, the single hotspot location specific to the patient was documented.

After identifying the individual motor hotspots using TMS, the EEG data were sampled at a rate of 500 Hz using a multi-channel EEG acquisition system (Liveamp, Brain Products GmbH, Gilching, Germany). The twenty-nine EEG electrodes were attached to the scalp using the international 10–20 system. The ground and reference electrodes were attached to Fpz and FCz, respectively (Figure 4(a)). The participants completed a simple hand movement task to measure EEG data. As introduced in ‘3. EEG Measruement’, the hand movement task, and rest were alternately performed for 30 trials for each hand. However, due to the limited motor ability of paralyzed patients, the finger-tapping task was replaced with a simple hand-grasp task. Thus, they lightly held their hand whenever a red circle appeared in the center of the monitor (Figure 4(b)). We excluded the EEGs of four subjects for the affected hand and three subjects for the unaffected hand due to the absence of motor hotspot lcoation. Additionally, we excluded the EEGs of three subjects for both hands due to recording errors and high contamination from physiological artifacts. Consequently, we used data from 23 and 21 subjects for the affected and unaffected hands, respectively, for data analysis.

**Figure 4.**
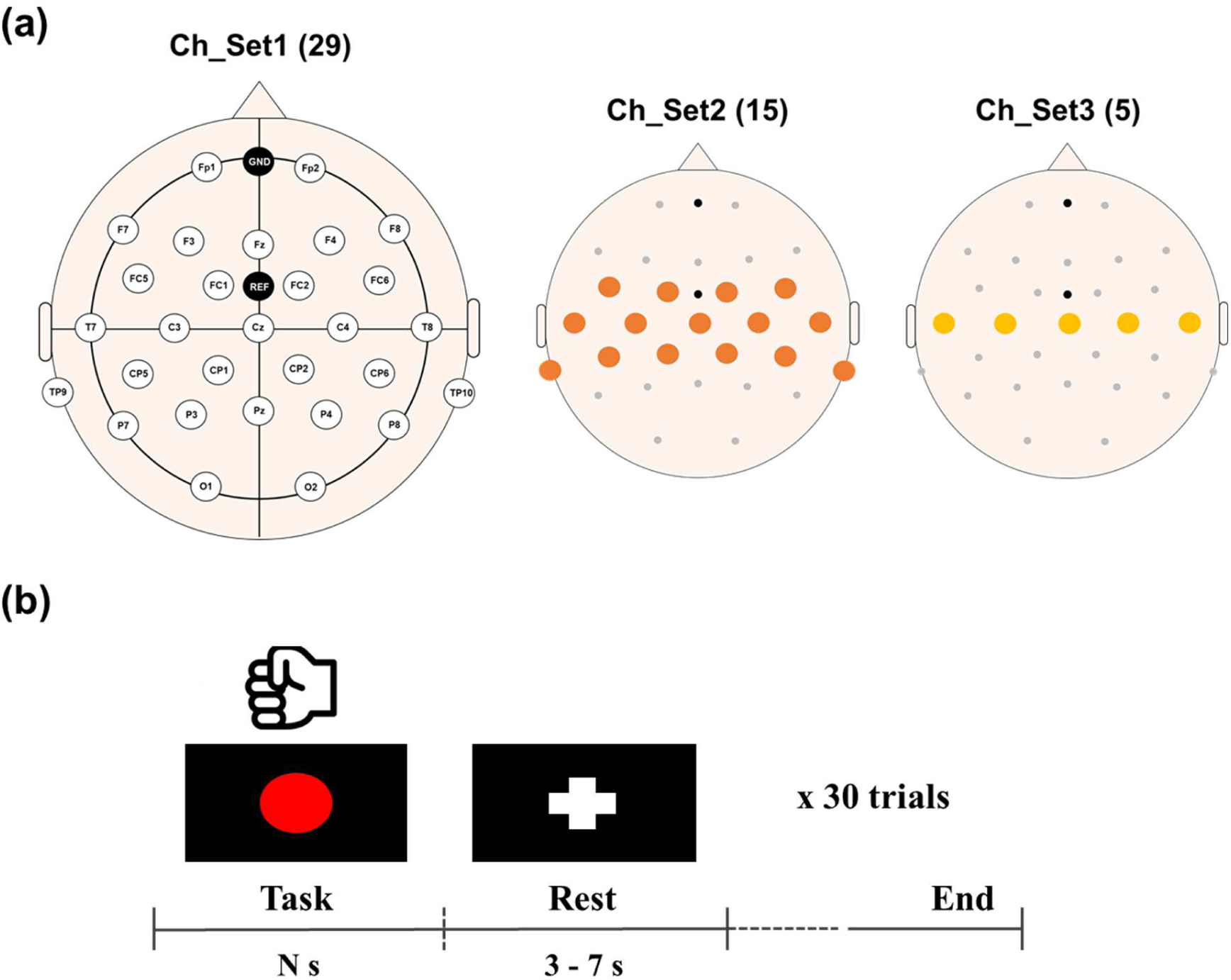
(a) Electrode attachment location used for EEG data in the stroke patient group. Three different channel sets are used for data analysis to determine the number of channels on the error distance in the motor hotspot location. The number in parentheses is presented as the number of channels in that channel set. (b) Experimental paradigm for stroke patient. Each patient performed the task of hand grasping within 2 seconds whenever a red circle appeared. At the end of the task period, the fixation (‘+’) mark is displayed to indicate a rest period.

The recorded data were analyzed following the procedures introduced in **‘4. Data Analysis’** and **‘5. Performance Verification’**. Exceptionally, the preprocessed data were segmented between -0.5 and 1 s whenever a red circle appeared in the center of the monitor during each trial. This segmentation procedure was applied because all patients completed the task within 1 s. In this analysis, the raw data (Input_1) were only used as input for identifying motor hotspots in stroke patients, as shown in Figure 6.

## Results

Figure 5 presents the topographic event-related spectral perturbation (ERSP) maps for 8 - 30 Hz of each hand movement task for each healthy and stroke group. The healthy group (Figure 5(a)) had focal dominant event-related desynchronization (ERD) patterns around the bilateral motor area. In contrast, the stroke group (Figure 5(b)) represented a relatively low amplitude and broad ERD patterns when performing the movement task using the hemiplegia side.

**Figure 5.**
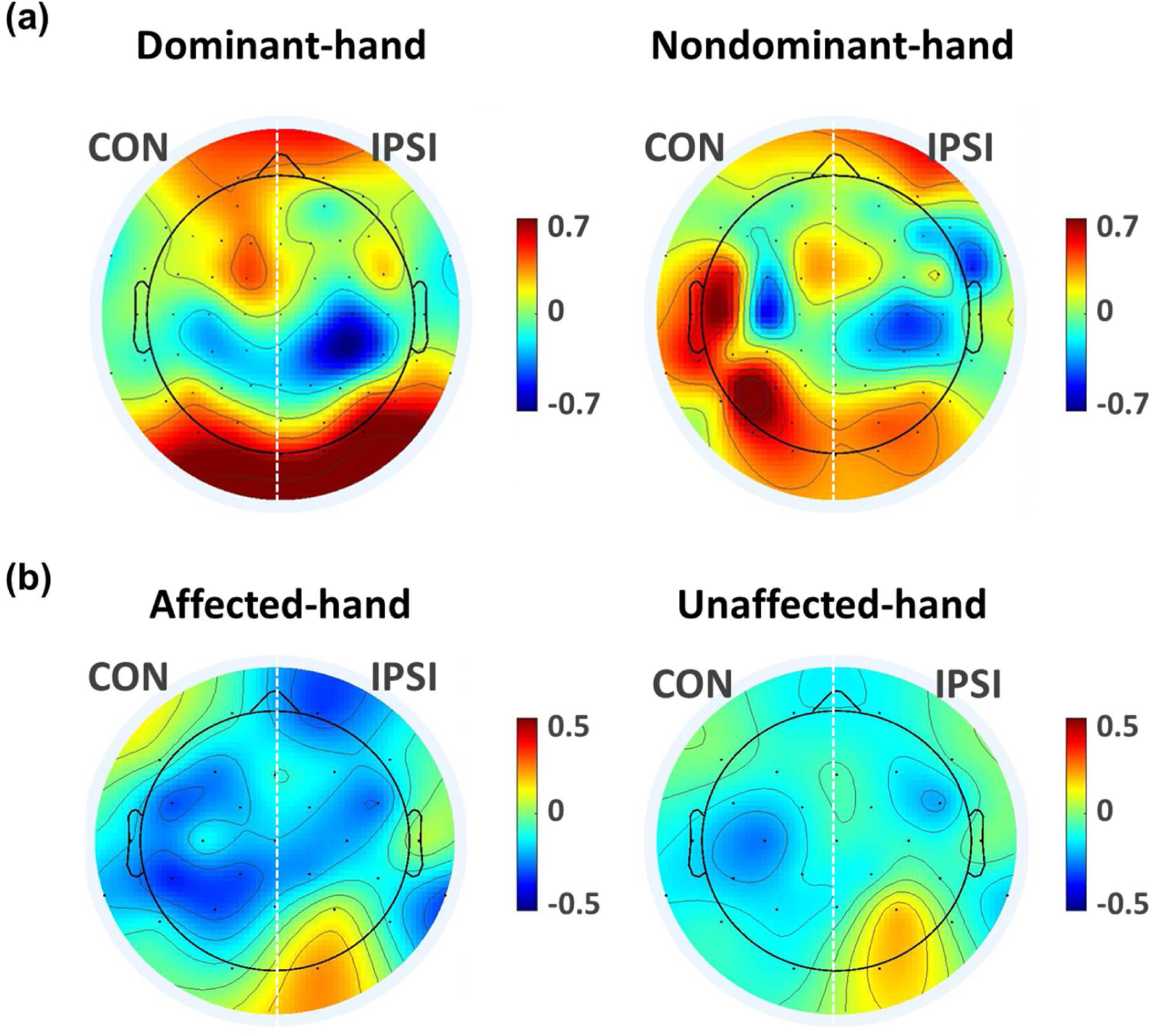
Grand averaged topographical event-related spectral perturbation (ERSP) maps for the alpha (8-13 Hz) and beta (13-30 Hz) bands associated with voluntary movements for each group. (a) Healthy group, (b) Stroke patient group. Although the strong ERD pattern appears around the motor area, the stroke group shows a relatively broad pattern when using the hemiplegia site. The terms CON and IPSI refer to the contralateral and the ipsilateral hemispheres, respectively, with respect to the hand being used.

Figure 6 shows the mean error distances of the estimated motor hotspot location using the EEG-based deep learning approach with 63 channels of healthy control. No significant difference was observed between the left and right hand for all conditions (two-sample t-test *p*-value < 0.05). The mean error distances consistently decreased as the raw data underwent increasingly advanced signal processing techniques informed by domain knowledge. However, even the input of raw data (Input_1) showed a low mean error distance of 2.27 ± 0.27 mm (RM-ANOVA, followed by paired-t-test for post hoc with Bonferroni corrected, *p*-value <0.05: mean error distances of each input type in order of Input_5 ≅ Input_4 ≅ Input_3 ≅ Input_2 < Input_1 for right-hand and Input_5 ≅ Input_4 ≅ Input_3 < Input_2 < Input_1 for left-hand). The commercial tES electrode size is typically at least 1 cm in diameter. Since the error distance of the motor hotspot identified by Input_1 (raw data) was considered acceptable, only the raw data were used for further analysis.

**Figure 6.**
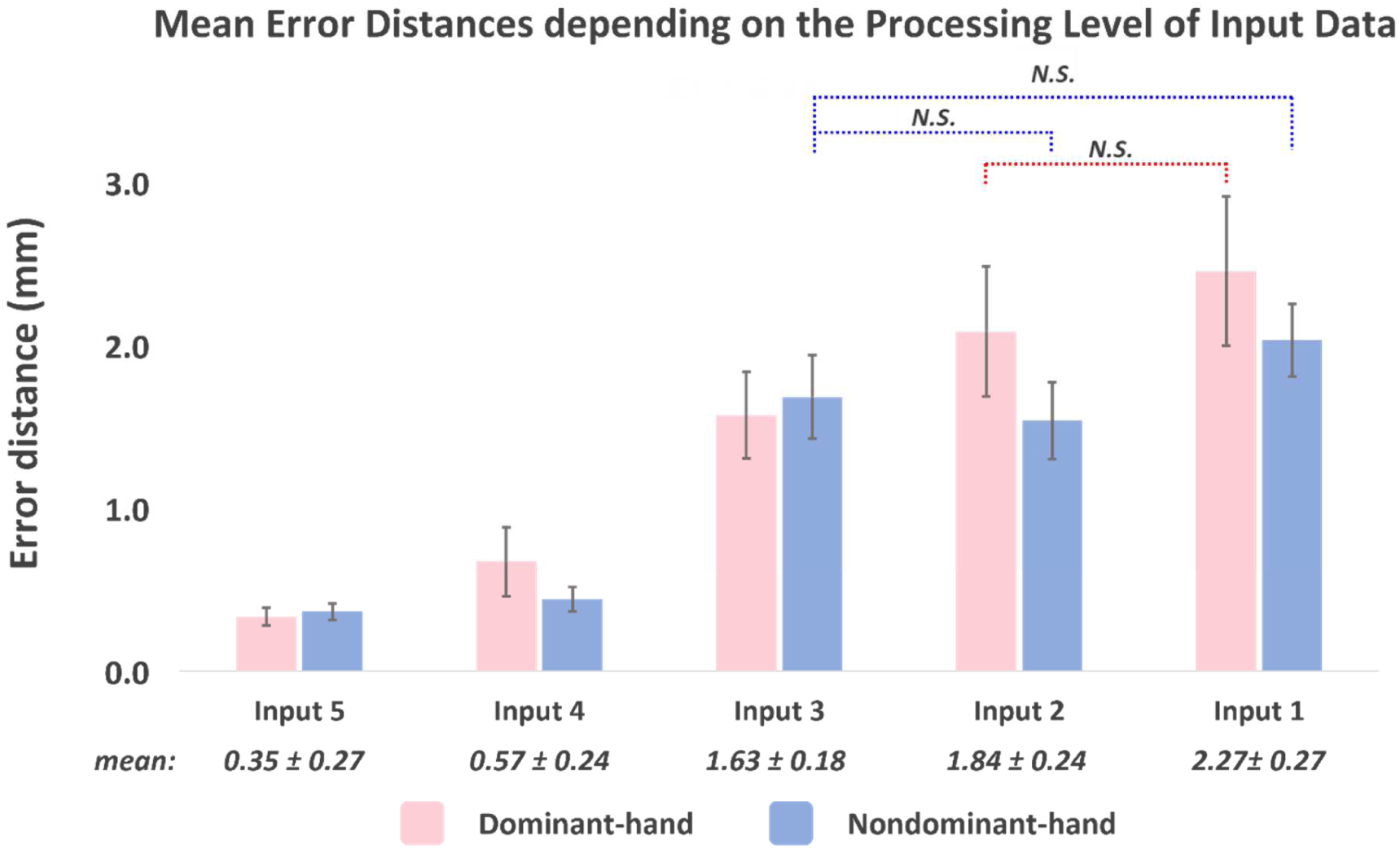
Mean error distances and standard errors between the motor hotspot locations identified by TMS-induced MEP and EEG-based depending on the processing level of input data (RM-ANOVA with Bonferroni corrected *p*-value < 0.05: Input 5 = Input 4 = Input 3 = Input 2 < Input 1 for right-hand and Input 5 = Input 4 = Input 3 < Input 2 < Input 1 for left-hand). The numbers below the bar graphs represent the mean error distances of the averaged both hands. No significant difference is observed between the left and right hand for all conditions in terms of the error distance (two sample t-test *p* > 0.05).

Figure 7 presents the mean error distances of estimated motor hotspot location for each hand, based on the number of channels using raw data (no significant difference is observed between the left and right hand for all condition, two-sample t-test, p-value < 0.05). The performance remained excellent, with a mean error distance of less than 1 cm in all conditions. Especially, the average error distance was 2.34 ± 0.19 mm, even when only using 9 channels around the motor area (RM-ANOVA, followed by paired-t-test for post hoc with Bonferroni corrected, *p*-value <0.05: mean error distances of each Ch_Set in order of Ch_Set1 ≅ Ch_Set2 ≅ Ch_Set3 ≅ Ch_Set5 < Ch_Set4 for right-hand and Ch_Set1 ≅ Ch_Set2 ≅ Ch_Set3 ≅ Ch_Set5 < Ch_Set4 for left-hand).

**Figure 7.**
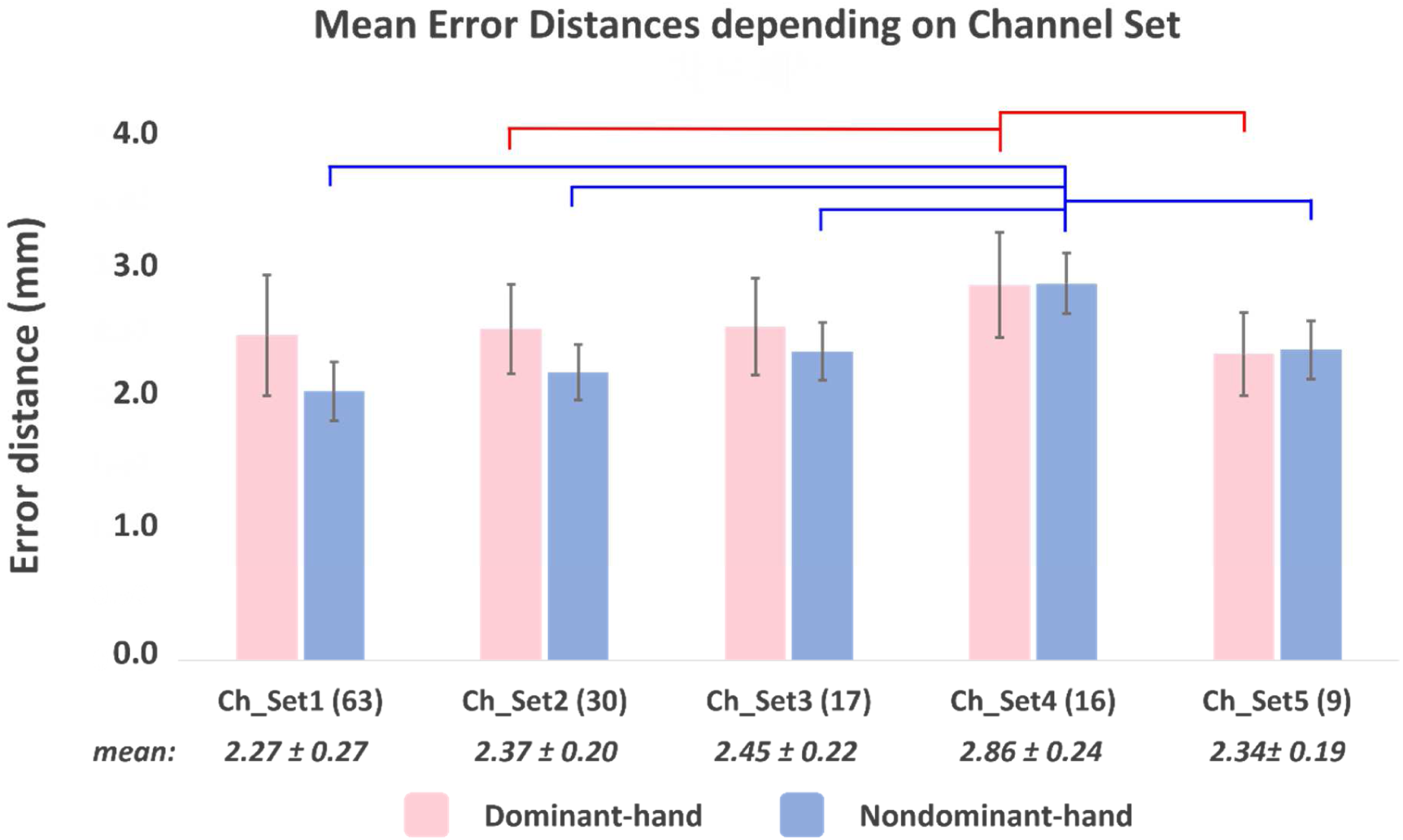
Mean error distances and standard errors between the motor hotspot locations identified by TMS-induced MEP and raw-EEG-based depending on the number of channels (RM-ANOVA with Bonferroni corrected *p*-value < 0.05: Ch_Set 1 = Ch_Set 2 = Ch_Set 3 = Ch_Set 5 < Ch_Set 4 for right-hand and Ch_Set 1 = Ch_Set 2 = Ch_Set 3 = Ch_Set 5 < Ch_Set 4 for left-hand). The numbers below the bar graphs represent the mean error distances of the averaged both hands. No significant difference is observed between the left and right hand for all channel sets in terms of the error distance (two-sample t-test *p* > 0.05). The solid lines indicate pairs with statistically significant differences.

Figure 8 presents the mean error distances of estimated motor hotspot location for each hand, based on the number of trials using 9 channels of raw data (no significant difference is observed between the left and right hand for all conditions, two-sample t-test, *p*-value < 0.05). Although the mean error distances increased as the number of trials used for analysis was reduced, it is important to note that the mean error distance remained below 1 cm even when the number of trials was reduced to 10 (paired t-test between the condition using 30 trials and the others, *p*-value < 0.05: mean error distances of each condition in order 30 ≅ 25 ≅ 20 ≅ 15 ≅ 10 < 5 for right-hand and 30 ≅ 25 ≅ 20 < 15 ≅ 10 < 5 for left-hand).

**Figure 8.**
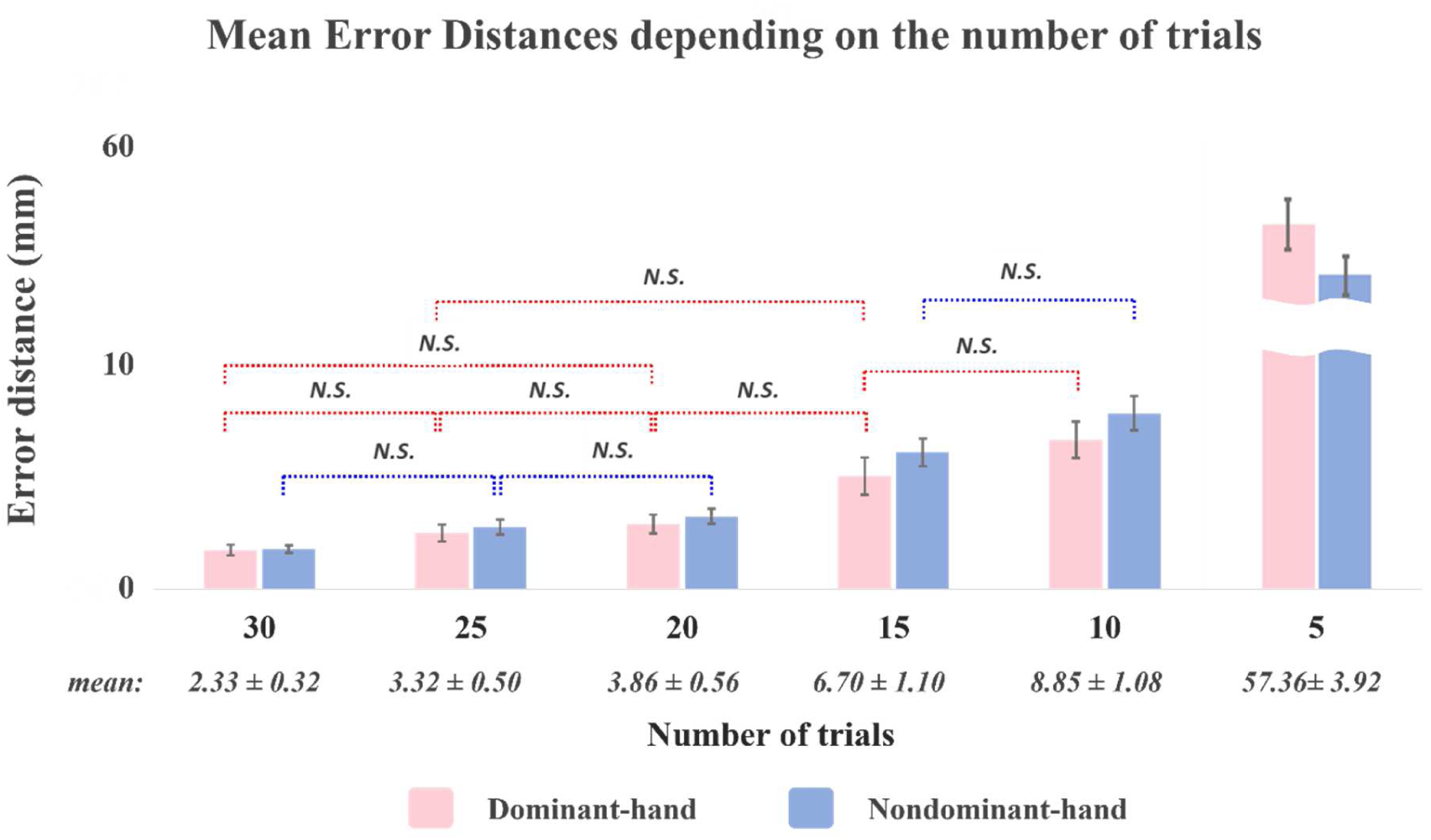
Mean error distances and standard errors between the motor hotspot locations identified by TMS-induced MEP and 9 channels of raw-EEG-based deep learning approach with respected to the number of trials (paired *t*-test between the condition using 30 trials and others, p-value < 0.05: mean error distances of each condition in order of 30 ≅ 25 ≅ 20 ≅ 15 ≅ 10 < 5 for right-hand and 30 ≅ 25 ≅ 20 < 15 ≅ 10 < 5 for left-hand). The numbers below the bar graphs represent the mean error distances of the averaged both hands. No significant difference is observed between the left and right hand for all channel sets in terms of the error distance (two-sample t-test, *p*-value > 0.05).

Figure 9 presents the mean error distances of estimated motor hotspot locations for each hand using raw data from stroke patients. No significant difference was found between the left and right hand for all conditions, as determined by a two-sample t-test with a *p*-value < 0.05. Despite the slightly different neurological patterns observed in the EEG of stroke patients compared to the healthy group (Figure 5), all channel sets had a consistent mean error distance of 1.64 ± 0.14 mm, regardless of the location of the lesion. In particular, the error distance was found to be only 1.77 ± 0.15 mm when using only 5 channels around the motor area (RM-ANOVA, followed by paired t-test for post hoc with Bonferroni corrected, *p*-value < 0.05: mean error distances of each Ch_Set in order of Ch_Set1 < Ch_Set2 ≅ Ch_Set3 for affected-hand and Ch_Set1 ≅ Ch_Set3 < Ch_Set2 for unaffected-hand).

**Figure 9.**
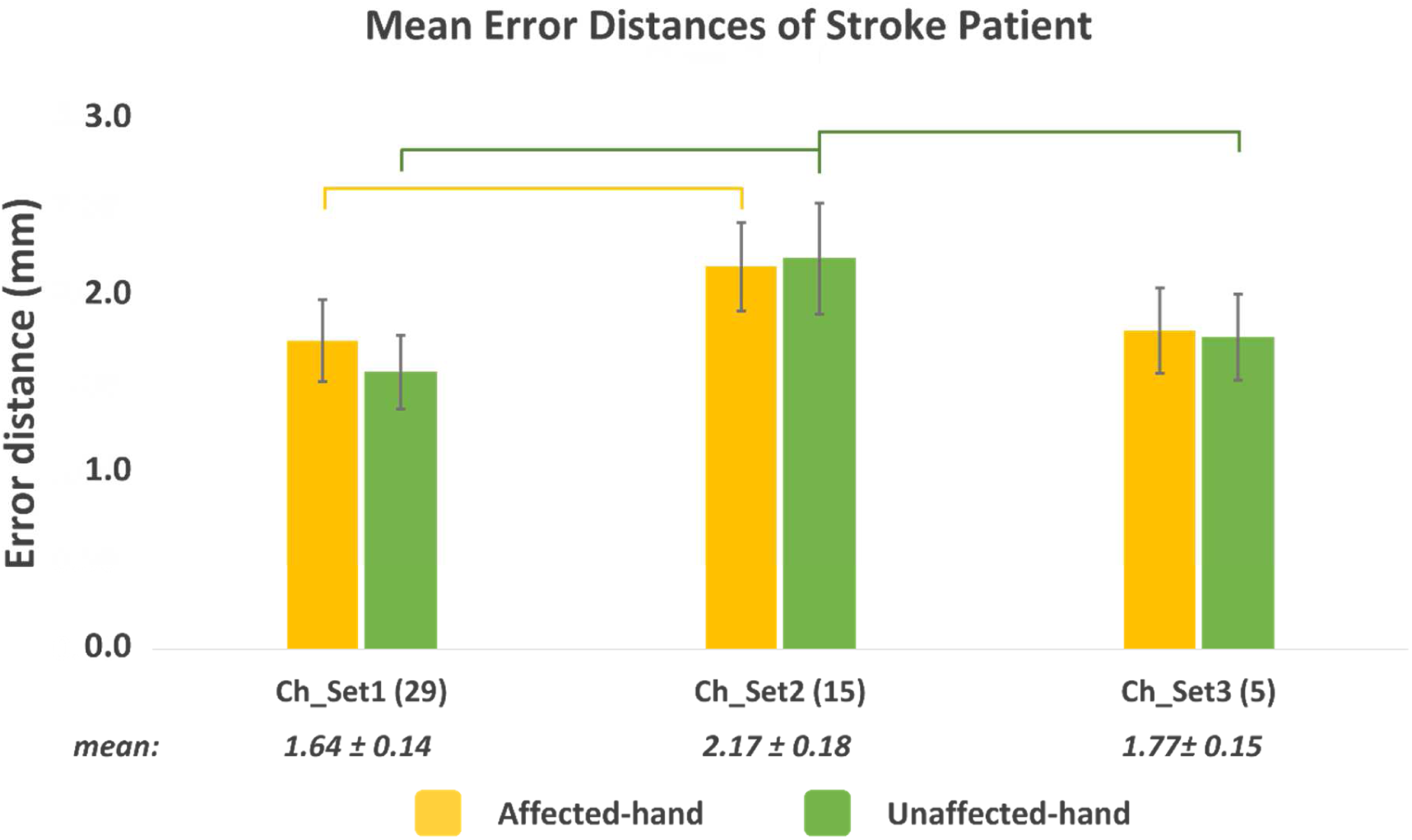
Mean error distances and standard errors between the motor hotspot locations identified by TMS-induced MEP and raw-EEG-based from the stroke patient depending on the number of channels (RM-ANOVA with Bonferroni corrected *p*-value < 0.05: Ch_Set 1 < Ch_Set 2 = Ch_Set 3 for affected-hand and Ch_Set 1 = Ch_Set 3 < Ch_Set 2 for unaffected-hand). The numbers below the bar graphs represent the mean error distances of the averaged both hands. No significant difference is observed between the affected and unaffected hand for all channel sets in terms of the error distance (two-sample t-test *p* > 0.05). The solid lines indicate pairs with statistically significant differences.

## Discussion

In this study, we proposed an advanced EEG-based motor hotspot identification algorithm using deep learning. This approach aims to replace the traditional motor hotspot identification method by using TMS. The mean error distance was 0.35 ± 0.27 mm when utilizing the PSD of gamma band, which was found to be the most effective feature with the lowest error distance in our previous study [37]. The performance of identifying motor hotspots using EEG-based methods has improved by approximately 82% compared to the previous ANN model (2.18 ± 0.26 mm).

We also evaluated the signal processing capability of our proposed deep learning model based on the motor hotspot identification performance by varying the handcrafted signal processing level of the input data. The mean error distance was 2.27 ± 0.27 mm when using the raw data (Input_1) as input. Although there was a statistical difference compared to using PSD features (Input_5) extracted by professional signal processing based on feature engineering, the predicted error distance was reasonable, considering the commercial tES electrode size is over 10 mm in diameter. Moreover, the FDI-MEP can be evoked at a maximum of about 18 mm from the center of the motor hotspot [42]. It demonstrates the exceptional signal processing and feature extraction performance of the proposed algorithm, substantiating its high usability.

In addition, we confirmed the change in the identification performance by reducing the number of channels and trials conducted for analysis, in order to improve the practicality of proposed model. The results indicated a slight quantitative difference regarding to the number of channels, but the identification performance remains robust. The study found that using only nine channels around the motor area resulted in a mean error distance of 2.34 ± 0.19 mm. This contrasts with the previous study’s ANN model, where the mean error distances increased linearly as the number of channels decreased. The practicality of the proposed algorithm is demonstrated that the time required for EEG electrode placement, based on the 10–20 system with 19 channels, can over 30 minutes, depending on the expertise of the experimenter. Additionally, the algorithm for identifying motor hotspots demonstrated robust performance in terms of trial numbers. Even when analyzed based on only 10 trials, the error distance consistently remained below 1 cm.

Finally, we confirmed the clinical benefit of proposed algorithm by analyzing EEGs from stroke patients. It is important to validate the proposed motor hotspot identification method with patients to demonstrate its clinical feasibility, as the EEG patterns of patients with motor impairments may differ from those of healthy individuals. As expected, the stroke group showed different neural activity patterns compared to the healthy group during voluntary movement (Figure 5). The use of the hemiplegia site showed a relatively weak and broad pattern [43–45]. Nevertheless, the proposed EEG-based motor hotspot identification algorithm demonstrated strong identification performance, with a mean error distance of less than 1.77 mm, even when using at least 5 channels around the motor area. Interestingly, the motor hotspot identification performance of stroke group showed similar levels of identification performance despite using fewer channels than the healthy control group. Further research is necessary to identify the underlying cause, but scalp-recorded EEG data reveals intricate signals that involve the electrical activity of various neurons. Therefore, it is expected that even a small number of channels can effectively capture information from neurons near the primary motor cortex.

The traditional approach to identifying motor hotspots using TMS usually takes at least 30 minutes or longer. This process relies on the subjective judgment of inspector. Additionally, the manual targeting of identified hotspot location for the application of tES poses challenges and may lead to inaccuracies. Taking this aspect into consideration, the integration of the developed algorithm into the commercialized portable tES-EEG devices (e.g., Starstim by Neuroelectrics in 2012, M×N-5 and M×N-9 HD-tES by Soterix Medical in 2014) enables the automatic identification of motor hotspots and the simultaneous application of tES without relying on empirical judgment. The proposed approach for identifying motor hotspots has the potential to enhance usability for patients with impaired mobility, thereby reducing the necessity for frequent hospital visits. Additionally, it provides economic benefits by replacing the need for a TMS device with a relatively economical EEG device.

Moreover, tES-based neurorehabilitation has been shown to induce changes in brain activity patterns through neuroplasticity [36, 46]. Hence, the performance of the proposed algorithm may deteriorate in a longitudinal neurorehabilitation scenario due to changes in EEGs caused by neuroplasticity. In addition, individual differences exist in the time it takes for rehabilitation effects to manifest and the extent of change in EEG patterns. By continuously fine-tuning the user’s novel EEG data using the transfer learning concept, the individualized model proposed in this study can prevent model performance degradation caused by intra-subject variability. This approach avoids the need for repeated model construction and ensures consistent performance over time. In this respect, the proposed approach for identifying motor hotspots using EEG is expected to facilitate home-based motor rehabilitation using tES techniques for patients with impaired mobility. While determining the initial motor hotspot location is necessary for constructing the model, the increasing practical applicability of tES-based rehabilitation is expected to enhance patient compliance and participation in neurorehabilitation. Consequently, this is expected to improve rehabilitation outcomes.

Furthermore, the tES is not limited to exercise rehabilitation. tES can be applied to cognitive function and alleviation of symptoms of mental illness [47–51]. Typically, tES electrodes for cognitive rehabilitation purposes are attached to F3 and F4 to stimulate the dorsolateral prefrontal cortex (DLPFC), which is known to be associated with various cognitive functions [52–53]. Nevertheless, similar to the individual variability in the preferred exercise hotspot, it can be inferred that the target location for tES in cognitive rehabilitation may also exhibit slight variations among individuals. Therefore, we believe that the proposed algorithm can be expanded to a variety of applications, including cognitive rehabilitation and alleviation of mental illness symptoms.

## Conclusion

In this study, we proposed the advanced EEG-based motor hotspot identification using a deep learning algorithm and confirmed the practicality of our proposed deep learning algorithm by changing the identification performance while reducing the number of channels and trials. Additionally, we demonstrated the clinical benefits of our algorithm by utilizing EEG data from stroke patients.

## Acknowledgements

This work was supported by the National Research Foundation of Korea(NRF) grant funded by the Korea government (MSIT) (No. RS-2023-00302489), by National Research Foundation of Korea (NRF) grant funded by the Korea government (MSIT) (No. NRF-2022R1A6A3A13053491), by the MSIT(Ministry of Science and ICT), Korea, under the ITRC (Information Technology Research Center) support program (IITP-2023-RS-2023-00258971) supervised by the IITP (Institute for Information & Communications Technology Planning & Evaluation), by National Research Foundation of Korea (NRF) grant funded by the Korean government (MSIT) (NRF-2022R1A2C1006046).

## Competing interests

GYC, NJP, WSK, and HJH have a patent-pending entitled “Apparatus and method for detecting individual motor hotspot position based on brainwave using convolutional neural network, and transcranial electrical stimulation apparatus using the same”, number 10-2020-0133831, number PCT/KR/2021/002118, and number J10178.0001.

## Notes

### Summary of Updates

We fixed the broken image during the upload process and addressed the competing of interest.

